# Conversational Chemistry: A Novel Approach to Chemical Search and Property Prediction

**DOI:** 10.1101/2023.11.11.566721

**Authors:** JJ Ben-Joseph, Tim Oates

**Affiliations:** University of Maryland, Baltimore County, TensorSpace Inc.; University of Maryland, Baltimore County

## Abstract

We have developed an approach to train a chemical property prediction model using both English and the SELFIES chemical language describing the structure of small, drug-like molecules. This model generates chemical embedding vectors, which we then use to train classification models. Our straightforward softmax classification model surpasses the commonly-used message passing neural network architecture in certain chemical property prediction tasks. Moreover, these chemical embedding vectors can be employed in other applications, such as building a chemical search engine that enables users to find new drugs with natural language queries (e.g., “low toxicity blood brain barrier permeable drug that inhibits HIV replication”).

## 1 Introduction and Related Work

The significance of health in our overall well-being and happiness is indisputable. With an annual revenue of $1.2 trillion, the pharmaceutical industry plays a critical role in discovering, developing, and producing new therapeutic drugs [1]. Even minor advancements in the drug discovery and development process can profoundly impact human health, with substantial social and economic consequences [2]. Chemical property analysis is also essential i[article218||byline||]n material sciences [3], with numerous applications in fields such as photovoltaics, battery science, manufacturing, and food science.

Small molecules, often the active ingredients in therapeutic drugs, are preferred due to their chemical stability, ease of transport and production, and suitability for pill formulation. A major challenge in drug development is toxicity, as highly reactive compounds tend to be toxic, while completely inert compounds lack on-target effects. This highlights the need for robust, chemically active molecules that do not interfere with the body’s normal functioning.

Machine learning can aid in screening molecules for drug development, helping to narrow down potential therapeutics by eliminating likely toxic candidates and reducing the need for costly in-vitro (lab) experiments. Additionally, it can optimize specific properties, such as blood-brain barrier penetration for brain-targeting drugs. Machine learning can also help select small molecules that appear to interact or disrupt a particular protein target. Accelerating the screening of new molecules, especially during early development stages, can significantly enhance the drug development process’s efficiency and facilitate the creation of better drugs more rapidly [2].

However, many crucial chemical properties cannot be calculated analytically. Currently, these properties are imperfectly determined through expensive in-vitro assays or occasionally animal models, which are not only costly but also raise ethical concerns. Moreover, drugs can fail in clinical trials due to excessive toxicity, as seen in numerous examples within the ClinTox dataset in the MoleculeNet database [4], an open dataset that tracks clinical trials. These challenges have prompted extensive research into in-silico methods, often employing machine learning, to determine chemical properties.

### 1.1 Data representations of molecules

One of the challenges in machine learning is determining how to convert data into a representation that is suitable for machine learning, storage, and analysis. A common in-silico method for representing small molecules is through the use of SMILES strings [5]. SMILES (Simplified Molecular-Input Line-Entry System) strings are a way of representing small molecules in a computer-readable format. They are human-readable sequences of characters that describe the atoms and bonds in a molecule. Each element in the molecule is represented by a specific symbol, and the bonds between the atoms are represented by a specific arrangement of characters. For example, the SMILES string for aspirin would be `CC(=O)OC1=CC=CC=C1C(=O)O` representing the atoms and bonds between them. Table 1 contains all the major SMILES symbols.

**Table 1:** SMILES is a graph description where symbols have chemical meanings.

| Bond Token | Description |
| --- | --- |
| C | Organic Elements (Carbon) |
| [Au] | Non-Organic Element (Gold) |
| . | Non-Bond |
| - | Single Bond |
| = | Double Bond |
| # | Triple Bond |
| \$ | Quadruple Bond |
| : | Aromatic Bond |
| / | Stereoisomer (Opposite) |
| \ | Stereoisomer (Same) |
| ( | Begin Group |
| ) | End Group |
| @@ | Tetrahedral Chirality (Clockwise) |
| @ | Tetrahedral Chirality (Counterclockwise) |

**Table 2:** MoleculeNet classification datasets used in our work.

| Name | # Examples | # Tasks | Description |
| --- | --- | --- | --- |
| Tox21 | 8014 | 12 | Multi-organ toxicity assays |
| BBBP | 2039 | 1 | Blood-brain barrier permeability |
| HIV | 41127 | 1 | Interferes with HIV replication |
| ToxCast | 8575 | 617 | In-vitro toxicity screening assays |
| SIDER | 1427 | 27 | Adverse drug reactions |
| ClinTox | 1478 | 2 | Failed clinical trials due to toxicity |
| BACE | 1513 | 1 | Inhibits BACE-1 (Alzheimer’s disease) |

SMILES strings are human-readable sequences of characters that describe the atoms and bonds in a molecule, but do not encode any semantic information about the chemical properties of the molecule. There are certain mechanistic properties of molecules, such as binding energies with other molecules, that may be of interest, but these properties are computationally expensive to calculate exactly [6] and are non-linear. It would be useful to be able to predict these properties with high accuracy using machine learning methods. Additionally, even SMILES strings that appear similar can have very different properties due to the non-linear nature of molecular dynamics, and it is possible to create chemically invalid SMILES strings [7], making them challenging to use in chemical generator models.

Despite these challenges, machine learning models can still learn useful models from SMILES strings [8]. This is because the model is able to learn useful parameters that allow it to separate the input sequences in a non-linear fashion. SMILES strings are sometimes fed directly into models that work with sequences, borrowed from natural language processing, or the graph representation of the molecule is generated and passed into models that work on graphs, such as graph convolutional neural networks.

Another method for representing small molecules is through the use of vector representations, often called molecular fingerprints. Molecular fingerprints play a crucial role in molecular property prediction for drug discovery and many other tasks [9]. They can be considered a fixed-length feature vector derived from a SMILES string, providing a more structured representation of the molecule. However, no universal representation is suitable for all tasks. For instance, peptides or genomic sequences, which are also considered molecules, are typically represented at a higher level of abstraction than atoms and bonds [10]. Additionally, quantum physical representations that include the positioning of atoms in 3D space can be computed from a molecule’s SMILES description [11]. Molecular fingerprints are created by applying fixed or analytical algorithms to the molecule, rather than directly passing atoms and bonds to the model. This exposes information to the model that might not be discovered through standard training procedures. In some cases, the molecular fingerprint may even include aspects of the target property that the model aims to predict as part of the input itself.

The Morgan fingerprint [12] is a widely cited and studied molecular fingerprint, boasting competitive performance on cheminformatics tasks [13]. It represents each atom in a molecule as a node in a graph, with the bonds between atoms depicted as edges. The fingerprint is generated by calculating the circular fingerprints of the graph, which are fixed-length bit vectors representing the local structure around each atom. These bit vectors are then combined to create the final Morgan fingerprint—a fixed-length vector representing the entire molecule.

Efforts have also been made to create fingerprints that combine information from multiple approaches, such as those extracting features from both small and large molecules and combining them into a single fingerprint [10]. Descriptastorus [14] is a tool that creates a fingerprint using 200 different descriptors from RDKit.

RDKit [15] is an open-source cheminformatics and machine learning software toolkit. It assists researchers and developers in working with small molecules and chemical data across various applications, including drug discovery, chemical analysis, and data analysis. RDKit offers a wide range of capabilities, such as generating 2D and 3D chemical structures, calculating chemical properties and descriptors, performing chemical reactions, and applying machine learning models to chemical data. It is widely used in the pharmaceutical and chemical industries, as well as in academia, and is available under the BSD license. RDKit descriptors are functions that take a molecule as input and return a scalar value of interest, like molecular weight or solubility. By utilizing a fixed set of 200 of these descriptors, Descriptastorus can create a useful fingerprint for small molecules.

### 1.2 Message passing and graph convolutional models with molecular fingerprints

Using a 2D graph to represent a molecule is a popular approach for handling chemical data in machine learning applications. In this representation, atoms in the molecule serve as nodes in the graph, while bonds between atoms act as edges. This enables analysis of the molecule through graph algorithms and graph neural networks, specifically designed for graph-based data.

A key advantage of depicting molecules as 2D graphs lies in the ability to incorporate more information about the chemical structure beyond just atoms and bonds. For instance, node and edge features in the graph can encompass details about atomic elements, bond types, and other structural aspects of the molecule. This extra information can enhance the accuracy and performance of machine learning models applied to the data.

Another benefit of this representation is its capacity to intuitively and interpretably capture the connectivity and structural relationships within the molecule. This is particularly valuable for tasks like predicting chemical properties, where comprehending the structural foundation for the property is crucial.

The Message Passing Neural Network (MPNN) [16] is a highly successful approach that utilizes the 2D graph representation of molecules. In this method, edges in a graph are treated as paths for activations, or “messages,” to flow between nodes. This allows the model to capture essential information about the relationships between atoms and bonds in the molecule, as well as any additional features included in the graph representation.

MPNN operates by iteratively updating the node features through a series of message-passing steps. In each step, nodes receive messages from their neighbors, which are then combined and processed to update the node features. The updated features capture the local information and structural context around each atom, enabling the model to learn complex patterns in the molecular structure. After a fixed number of message-passing steps, the final node features are aggregated to produce a fixedsize representation of the entire molecule, which can then be used for downstream tasks such as property prediction. Our work builds upon the chemprop open-source toolkit, which was developed to showcase the capabilities of the MPNN, and we compare our results to this model.

MoleculeNet [4] serves as a widely utilized dataset for assessing the performance of diverse models in relation to chemical property tasks, such as toxicity and activity against biological targets linked to diseases. The integration of MPNNs and Descriptastorus-style fingerprints has demonstrated exceptional performance, achieving state-of-the-art or close to state-of-the-art results on multiple tasks within the MoleculeNet dataset.

### 1.3 Natural language processing and embedding models

Natural language processing (NLP) models have experienced significant advancements in recent years, thanks to being trained on extensive amounts of unlabeled text data. Transfer learning, a technique that fine-tunes these models using smaller data quantities for specific tasks like Named Entity Recognition (NER), has played a crucial role in this progress.

The concept of employing joint vector spaces to represent different data types, such as natural language and images, has a well-established research history. One notable example is the CLIP model [17], a deep neural network jointly trained on natural language and images. CLIP generates embedding vectors, which are fixed-sized representations of images or text that can be used directly in a classifier or for vector search tasks. For instance, integrating CLIP with a nearest neighbor search can develop an image search engine that responds to natural language queries. This is because the vectors for phrases like “cat sitting on a desk” and images of cats sitting on desks tend to be close together in geometric space when processed through CLIP.

The CLIP model creates a shared vector space by utilizing a contrastive loss function that encourages similar data points to be near each other in the vector space, while pushing dissimilar data points further apart. The model is trained on large amounts of data from various sources, including text and images from the internet. During this process, CLIP learns to create embedding vectors for each data point, capturing the underlying features of the data in fixed-sized representations.

Upon training completion, the model can generate embedding vectors for new data points. These vectors can be used directly in a classifier or for vector search tasks, such as image search using natural language queries. CLIP has proven effective in producing high-quality embedding vectors that capture relationships between different data types and can be applied to a wide range of tasks.

### 1.4 Transformer models and Large Language Models (LLMs)

Transformer models have become a significant breakthrough in natural language processing and machine learning. Introduced by Vaswani et al. [18], the transformer model is an architecture designed to overcome the limitations of recurrent neural networks (RNNs) and convolutional neural networks (CNNs) in processing sequential data. The transformer model relies on self-attention mechanisms, enabling it to capture long-range dependencies in sequences and process data in parallel, leading to better performance and faster training times compared to RNNs and CNNs.

A key feature of transformer models is their ability to scale with larger amounts of data and model parameters. This scaling capability has given rise to Large Language Models (LLMs), which are transformer models trained on massive text corpora, resulting in improved language understanding and generation capabilities. Some well-known LLMs include GPT-3 [**?**] and BERT [**?**], which have achieved state-of-the-art performance on various NLP tasks such as machine translation, sentiment analysis, and question-answering.

LLMs benefit from a pretraining-finetuning paradigm. In the pretraining phase, models are trained on vast amounts of unlabeled text data using unsupervised learning objectives, such as masked language modeling for BERT or autoregressive language modeling for GPT-3. This pretraining allows LLMs to learn general language features, including syntactic structures and semantic relations, by capturing the statistical properties of the training data.

In the finetuning phase, LLMs are adapted to specific tasks using smaller labeled datasets. This transfer learning approach enables the models to leverage their pretrained knowledge to achieve high performance on various tasks, even with limited labeled data. It has led to state-of-the-art results across a wide range of NLP benchmarks, such as GLUE [**?**] and SuperGLUE [**?**].

Despite their success, LLMs also come with certain challenges, such as their large size, which can make them difficult to deploy in real-world applications or on resource-constrained devices. Moreover, they can sometimes generate text that may be biased or offensive, as they learn from the data available on the internet, which can contain biased or controversial content. Ongoing research is addressing these challenges by developing techniques for model compression, fine-grained control over text generation, and mitigating biases in the training data.

#### 1.4.1 MiniLM Transformer

The transformer model is a neural network architecture that was developed by researchers at Google for natural language processing (NLP) tasks, such as language translation and language modeling. It has since been applied to a wide range of tasks in NLP and other fields, such as speech recognition and image generation.

The transformer model is based on the idea of using self-attention mechanisms to process input data in a parallel manner and capture long-range dependencies between input elements. It consists of encoder and decoder modules, which are connected by a multi-headed attention mechanism.

To process input data, the transformer model first encodes the input sequence into a sequence of hidden states using the encoder module. It then uses the attention mechanism to calculate the relationships between different elements of the input sequence, allowing it to capture long-range dependencies and contextual information. The decoder module then uses the hidden states and the attention mechanism to generate the output sequence.

The MiniLM transformer model is a small version of the transformer architecture that was developed by researchers at Google. It is a neural network model that is designed to process sequential data, such as natural language text or audio data, and perform tasks such as language translation or text generation.

The transformer architecture is based on self-attention mechanisms, which allow the model to process input sequences in a parallel manner and capture long-range dependencies between input elements. The MiniLM transformer model is a compact version of the transformer architecture that uses a smaller number of parameters and has a simpler structure, making it more efficient to train and deploy.

To process input data, the MiniLM transformer model first encodes the input sequence into a sequence of hidden states. It then uses self-attention mechanisms to calculate the relationships between different elements of the input sequence, allowing it to capture long-range dependencies and contextual information. The model then decodes the hidden states to generate the output sequence.

## 2 Approach

Our approach involves training a NLP model on the joint distribution of natural language and language-like chemical descriptions. We have tried many variations of approach which we describe in the following section. We found this idea of joint training, while easy to describe, is not as trivial as it may seem. Doing it in a naive fashion will lead to a model that will not converge. What we describe is the approach we used to not only train this way successfully, but to a create a high performance model using this technique.

We test our embedding model on the MoleculeNet dataset which is a benchmark of very practical and important tasks in drug discovery [19]. Before any further testing is done on a molecule, we must test the toxicity. We can not give a potentially toxic compound to humans in a clinical trial for instance, and overly toxic compounds are less useful in therapeutics. Several of MoleculeNet’s properties relate to toxicity. Notably it is one that can not be computed at all analytically, because it is not even a property of the molecule but rather the molecule and its interactions with the complex human metabolism. Other tasks include BBBP which classifies if a molecule can pass the blood-brain barrier, chemical properties related to the effect on HIV replication, as well as bioactivity against a Alzheimer’s disease target.

### 2.1 Dataset

We use MoleculeNet for both training and evaluation [4]. MoleculeNet is a large scale benchmark dataset for chemical property prediction using machine learning. MoleculeNet contains several public datasets and establishes standard metrics for evaluation.

We trained our model on three of the datasets in of MoleculeNet (Tox21, BBBP, and HIV). Since our model is not a classification model but rather an embedding model we decided to test our approach on the five datasets in this example without ever training our embedding on them at all.

MoleculeNet is organized in tasks which enable effectively mutliclass and multilabel classification. However, we are trying to map sentence pairs to each other in our work. So we created a dataset of chemical property descriptions written as sentences in the English language. We mapped these in pairs to chemical sequences in variations depending on the task at hand.

- **Tox21:** For this dataset, we took all 12 tasks and created a sum based on if they failed the toxicity assay. This created a mapping between each chemical in the dataset and a number between 0 and 9. We created an index of statements such as as seen in Table 3, where the number of failed assays is the index.
- **BBBP:** Since this had one task, we simply mapped the phrases *Does not enter the brain* and *Not a blood-brain permeable therapeutic drug* to not-permeable and *Drug that crosses the blood brain barrier* and *Blood-brain permeable therapeutic drug* to permeable.
- **HIV:** Similarly since this had one task, we mapped the phrases *Not a HIV therapeutic drug* and *Not a AIDS therapeutic drug* to no activity against HIV and *HIV drug therapeutic drug* and *AIDS drug therapeutic drug* to activity against HIV.

**Table 3:** Tox21 failed assays were summed for each compound and assigned a label as such.

| Mapped phrase | Number of failed toxicity assays |
| --- | --- |
| Safe therapeutic drug | 0 |
| Mostly safe therapeutic drug | 1 |
| Somewhat safe therapeutic drug | 2 |
| Therapeutic drug with risks | 3 |
| Chemical | 4 |
| Semitoxic chemical | 5 |
| Toxic chemical | 6 |
| Poison | 7 |
| Extremely poisonous compound | 8 |
| Very deadly poison | 9 |

**Table 4:** Our model trained with different sampling losses on Tox21 dataset of MoleculeNet showed that our novel loss function performed the best.

| Loss Function | Spearman correlation<br>(higher is better) |
| --- | --- |
| No Negative Sampling | 0.0212 |
| Binary (Triplet-like) Loss | 0.9130 |
| Pseudo Normal-Negative Loss | **0.9890** |

**Table 5:** The choice of molecular representation showed no statistical significance. Our final model used SMILES.

| Molecular Representation | Spearman correlation<br>(higher is better) |
| --- | --- |
| SELFIES | 0.9887 |
| SMILES | 0.9890 |

**Table 6:** The choice of base model showed no statistical significance. Our final model ended up using the simplest model all-MiniLM-L6-v2.

| Transformer Base Model | Spearman correlation<br>(higher is better) |
| --- | --- |
| all-mpnet-base-v2 | 0.9885 |
| all-distilroberta-v1 | 0.9888 |
| all-MiniLM-L12-v2 | 0.9875 |
| all-MiniLM-L6-v2 | 0.9890 |

### 2.2 Hardware setup

The experiments ran on our internal GPU machine containing four NVIDIA Tesla V100 GPUs (5120 computational cores over 84 SMs, 16 GB onboard memory) all connected via NVLink, two 18-core Intel Skylake CPUs and 384 GB of DDR4 memory. The experiments took about one week of usage on this machine to finish. The machine runs CentOS 7.9.

### 2.3 Using Pseudo Normal-Negative Loss with Negative Sampling

When we first started this research we only trained with positive examples. For example, a “highly toxic compound” was mapped to the chemical description of a highly toxic compound. This approach did not work. The models we trained using only positive sampling did not converge to any useful embedding at all.

What we did after was utilize the concept of negative sampling. Negative sampling is widely used in embedding model training. The FaceNet neural network [20] for face recognition used the concept of triplet loss, a positive example is trained with a positive anchor example and a negative example jointly.

Although a triplet-like loss did improve the performance significantly, we developed a new kind of loss that further improved the performance of the embedding model. We call this pseudo normal-negative loss because as opposed to the function being a true probability distribution, it pushes normal curve maximum close to 1.

First we start with a normal distribution *N* (*μ* = 0, *σ* = 2):

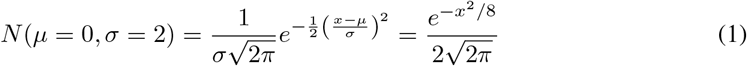

We further compute a scale factor *γ*. In this case 0.999 is the true positive value and the scaled top of the normal curve. We found that setting this very close to 1 but not exactly 1 was needed for the model to converge properly.

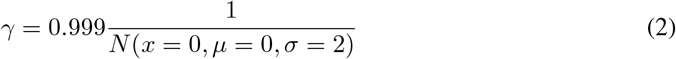

For each negative sample we compute a label as such, with 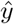 as the label, y as the ground truth, and i as the index of the prompt (Number of toxicity assays) as shown in table 3.

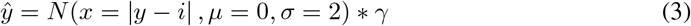

For positive examples this will return 0.999 but negative samples return lower numbers as part of normal curve.

### 2.4 Molecular Representation: SELFIES or SMILES?

SMILES is not the only chemical language in use in the field of cheminformatics. Some were developed especially to be useful in machine learning. The SELFIES language is one very notable example [7]. SELFIES differs from SMILES in that every possible SELFIES string describes a valid molecule. In SMILES it is very easy to create strings which contain syntax errors or chemically impossible molecules. This is not the case in SELFIES.

Our original work used SELFIES exclusively. This is because we encountered results in past unpublished work that showed that SELFIES also improves the performance of input-only models. Although using SELFIES did not harm our performance, we did not notice a difference in performance when used on our task. This can be explained by the fact that we do not output molecules, we convert inputs into embeddings. We have full control over the SMILES string inputs and thus all our inputs are a priori valid. Why SELFIES performed better on a similar input controlled research question we encountered is not yet understood.

However, SELFIES has advantages in being easier to parse and better suited for generative models. Further we developed our own SMILES-aware parser that converts SMILES strings into sensible word-like tokens. If we did not do this, it is possible that SELFIES which are already rather semantic and create easier to parse tokens would produce better results.

### 2.5 Choice of base NLP model

The choice of what NLP model to fine tune was explored early on. We tested with four different Transformer-based or inspired NLP models and we found no significant gains in performance based on which was chosen. For these results, each of these models were trained for 65 epochs on the Tox21 dataset alone.

- **all-mpnet-base-v2**: Based on MPNet [21], which combines permuted language modeling and also takes auxiliary position information as input which allows the model to see a full sentence thus reducing the position discrepancy.
- **all-distilroberta-v1**: The DistilBERT [22] model applied to Roberta. DistilBERT uses a technique called model distillation to reduce the size of BERT by as much as 40% while still being within 97% accuracy on the tasks tested.
- **all-MiniLM-L12-v2**: MiniLM [23] uses a procedure called deep self-attention distillation. The small model called a student is trained mimicking the self-attention module of the larger model (teacher).
- **all-MiniLM-L6-v2**: Same as above but with six layers instead of twelve. This is the actual platform that our model (as tested in the Results section) used because of its high computational performance compared to the others.

## 3 Results

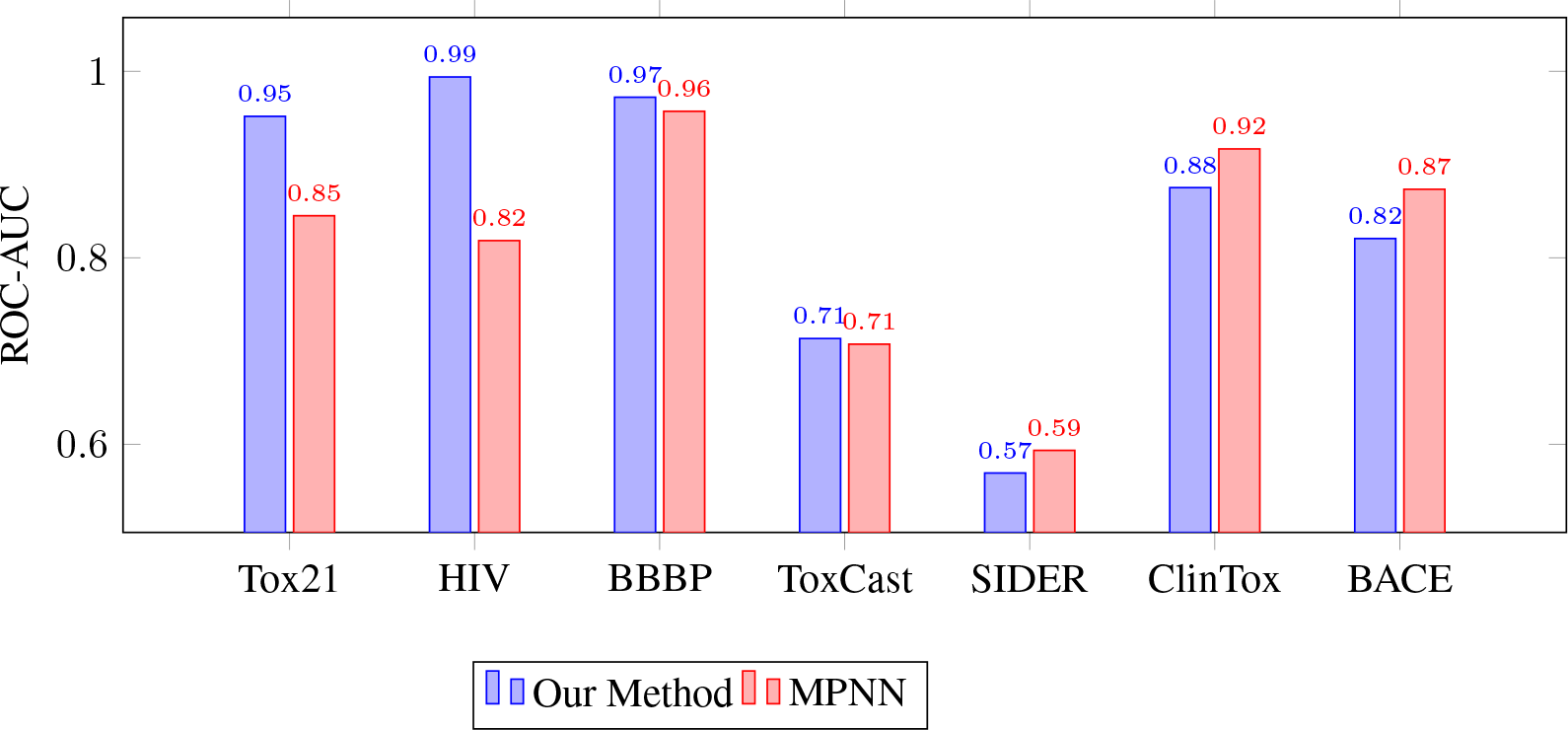

We ran experiments to compare the result of our model and the Message Passing Neural Network model (MPNN) on a selected number of MoleculeNet classification datasets described in the Approach section. Our approach used the standard hyperparameters of the MPNN in the chemprop toolkit (commit: b1f342255f26d40b65c34f1260297d2b772b98b2), as well as standard hyperparameters of the NLP models from Sentence-Transformers 2.2.0 [24]. The results of the embeddings are directly classified by a softmax output layer in both cases. We use the standard test/test/val split of 0.8/0.1/0.1 for both tests. The classification models of MPNN and our approach were both trained for 30 epochs utilizing chemprop.

The found the performance results to be rather intriguing. We trained our model on only three of the datasets in MoleculeNet: Tox21, HIV, and BBBP. And one can see obviously that the model does excellent on these datasets. We do use the same train/test/val splitting procedure in both. But in any case, at the minimum this means the model is in fact fitting the data, something we couldn’t accomplish without our negative sampling - it simply didn’t work. And our loss function improved things significantly.

Another intriguing result in our opinion is the result on the ToxCast dataset. Our model was not trained on it at all. Yet it performs slightly better compared to the well established MPNN model. Probably there is some kind of knowledge being transferred from the Tox21 data, because they are similar tasks. These are benchmarks that come from the same NIH lab, but they are different datasets with different tasks. The other datasets we tested does not advance past MPNN, however, it’s rather competitive. These are problems that our model has no understanding of and has not been trained on at all. If there is anything it is learning from the other datasets it is just the method of which to separate molecules and their general representation in Transformer form.

## 4 Drug discovery as large scale search problem

Our experiments focused on classification simply because classification provides the easiest results to interpret from a scientific perspective. But the true power in our approach stems from the fact that our model is an embedding model. In this section we show the possibility discovering new therapeutic drugs through search.

To prove this out we developed a rudimentary user interface that allows interactive queries against our model. For example a user could write a query such as *Low toxicity HIV therapeutic drug that crosses the blood-brain barrier*.

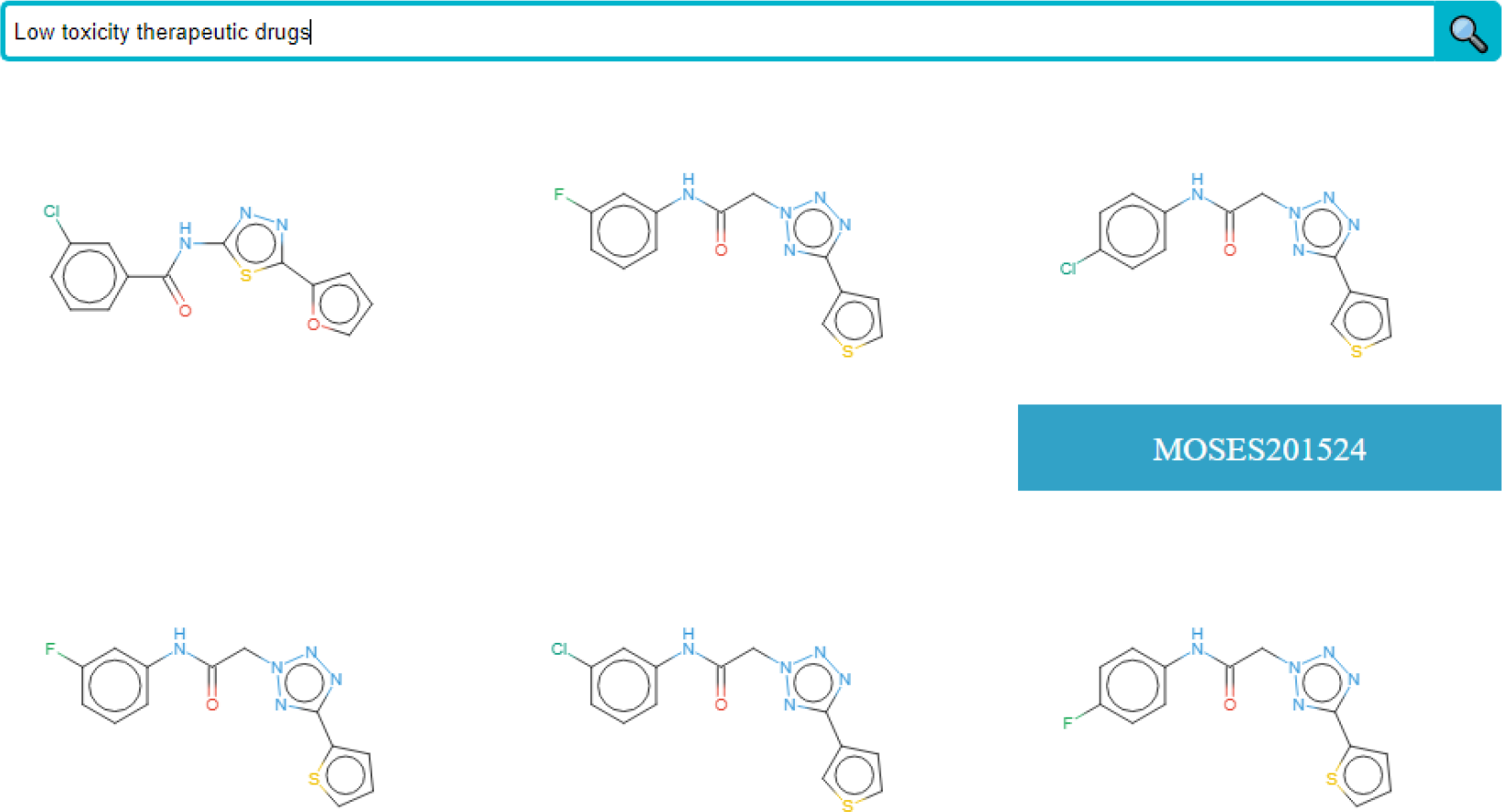

This whole UI and inference model runs on a AWS t3.micro instance with a swapfile. The results are from searching 1,250,000 of the MOSES dataset [25], a open source unlabeled dataset (MIT licensed) of drug-like molecules that could be manufactured. In the figure above you can see the tag assigned to each molecule when your cursor is hovered over it, and clicking on it produces the SMILES string. These results take a second or two at most to return. Our system uses the NGT toolkit and ONNG algorithm [26] to do fast approximate nearest neighbor search.

## 5 Limitations and Negative Social Impacts

This work was only trained on limited tasks of toxicity, blood-brain barrier permeability, and inhibiting HIV. Although this is a diverse set of tasks, it does not remotely capture all chemical properties of interest, and the dataset sizes are rather small especially when compared to what is available in NLP or computer vision. To address this, future work should train on more data, perhaps even scraping SMILES strings from clinical trials and using surrounding text as a prompt - a similar approach to CLIP. There will still be way less available data compared to images and text, a common combination all over the web.

This work could also be simply reversed to create the opposite of therapeutic drugs. For example instead of searching for drugs with low toxicity one could also look for drugs of maximum toxicity. Drug discovery systems are well known for having this dual-use problem, for example drug discovery systems have been used for discovering nerve agents [27].

## 6 Future Work and Conclusion

There are way more investigations we or others can pursue using this approach. Future work will include training our model with more datasets. An area of investigation we are very interested in is scraping academic research articles for SMILES strings and using the surrounding text or captions as input. This is a training method that can not be trivially accomplished with the structured classification models in use today and is similar to the training approach successfully used in CLIP. Another investigation worth trying is to use the approach in a generative fashion. To create molecules from textual descriptions. In these generative models we believe the SELFIES representation will come to play.

We showed that a NLP model with a novel training procedure can be competitive with a neural network architecture that is designed for the chemical property prediction task. Further we showed that our approach can generalize across tasks that it has been never trained on, learning some kind of abstract knowledge of the chemical space. Lastly we showed the potential of embedding models to solve new kinds of problems that are not possible with classification models like creating a search engine allowing a user to search for molecular properties using natural language.

